# Dynamic evolution of the MutS family in animals: multiple losses of MSH paralogues and gain of a viral MutS homologue in octocorals

**DOI:** 10.1101/2020.12.22.424024

**Authors:** Viraj Muthye, Dennis V. Lavrov

## Abstract

MutS is a key component of the Mismatch Repair (MMR) pathway. Members of the MutS family of proteins are present in bacteria, archaea, eukaryotes, and viruses. Six MutS homologues (MSH1-6), have been identified in yeast, three of which function in nuclear MMR, while MSH1 has been associated with mitochondrial DNA repair. MSH1 is believed to be lacking in animals, potentially reflecting the loss of MMR in animal mitochondria, and correlated with higher rates of mitochondrial sequence evolution. An intriguing exception has been found in octocorals, a group of marine animals from phylum Cnidaria, which encode a MutS-homologue (mtMutS) in their mitochondrial genome. It has been suggested that this protein functions in mitochondrial DNA repair, which would explain some of the lowest rates of mitochondrial sequence evolution observed in this group. To place the acquisition of mtMutS in a functional context, we investigated the evolution of the whole MutS family in animals. Our study confirmed the acquisition of octocoral mtMutS by horizontal gene transfer from a giant virus. Surprisingly, we found orthologues of yeast MSH1 in all hexacorals (the sister group of octocorals) and several sponges and placozoans. By contrast, MSH1 orthologues were lacking in octocorals, medusozoan cnidarians, ctenophores, and bilaterian animals. Furthermore, while we were able to identify MSH2 and MSH6 in all animals, MSH4, MSH5, and, especially, MSH3 were missing in multiple species. Overall, our analysis reveals a dynamic evolution of MSH family in animals, with multiple losses of MSH1, MSH3, some losses of MSH4 and MSH5, and a gain of octocoral mtMutS.

## 1 Introduction

The Mismatch Repair Pathway (MMR)^1^ is one of the five major pathways involved in DNA repair [1, 2, 3]. MMR corrects DNA mismatches generated during DNA replication, improving the fidelity of DNA replication by 50-to 1,000-fold [4]. Deactivation of this pathway in mammalian cells has been associated with predisposition to various types of cancer [5], defects in meiosis [6], and sterility [7] (summarized in [3, 8]). In organisms, like *Helicobacter pylori,* where MMR proteins are absent, loss of large genomic regions and elevated mutation rates have been reported [2, 9, 10].

MMR has been well-characterized in several prokaryotes and eukaryotes [11, 12]. The key proteins of MMR, MutS and MutL, are well-conserved in all three domains of life [13]. In *Escherichia coli,* a MutS homodimer recognizes and binds to the DNA mismatch. MutL interacts with MutS, and recruits and activates the endonuclease MutH. MutH introduces a nick in the newly-synthesized DNA strand that contains the error, which is followed by excision and re-synthesis of the error-containing strand [8]. The key aspects of bacterial MMR are conserved in eukaryotic MMR (DNA mismatch identification, error strand discrimination, excision, and DNA re-synthesis) [8]. However, the underlying machinery is more complex. At least nine MutS homologs have been identified in eukaryotes (Table 1). Of these, MSH2-6 are well-conserved in most eukaryotes, and are the only nuclear MutS homologues identified in animals. Unlike a single MutS homodimer in prokaryotes, two MutS heterodimers usually function in DNA mismatch recognition in eukaryotes: MutS*α* (MSH2 and MSH6) and MutS*β* (MSH2 and MSH3) [14, 8].

**Table 1:** Distribution and localization of MutS homologues in animals, plants, and fungi. The subcellular localization of the animal mitochondria-encoded MutS homologue (mtMutS) is not known.

| **Animals** | **Fungi** | **Plants** | **Genome** | **Localization** |
| --- | --- | --- | --- | --- |
| - | MSH1 | - | Nuclear | Mitochondria |
| - | - | Plant-MSH1 | Nuclear | Mitochondria<br>Chloroplast |
| MSH2 |  |  | Nuclear | Nucleus |
| MSH3 |  |  | Nuclear | Nucleus |
| MSH4 |  |  | Nuclear | Nucleus |
| MSH5 |  |  | Nuclear | Nucleus |
| MSH6 |  | Plant-MSH6<br>Plant-MSH7 | Nuclear | Nucleus |
| - | - | MutS2 | Nuclear | Nuclear |
| mtMuts | - | - | Mitochondrial | Mitochondria(?) |

While MMR in the nuclear genome has been well-characterized and extensively studied, organellar MMR is much less understood. Two MutS homologues in eukaryotes have been shown to function in organelles: fungal MSH1 and plant MSH1 (pMSH1)^2^. Even though both have been named “MSH1”, plant and fungal MSH1 are not orthologous. Yeast MSH1 is closely related to the rest of MutS homologues in the genome (MSH2-6). By contrast, most closely related homologues of pMSH1 are found in giant viruses [15]. A recent study showed that pMSH1 is dual-localized to mitochondria and chloroplast and involved in mismatch repair [15], while the role of yeast MSH1 in MMR remains controversial [16, 17, 18]. Within animals, the lack of a mitochondria-targeted MutS homologues suggests that organellar MMR may be absent. In fact, the lack of MMR in mitochondria has been reported as one of the causes of higher substitution rates in animal mitochondrial genomes compared to their nuclear counterparts [19]. However, our knowledge of DNA repair machinery in animal mitochondria is far from complete and mostly limited to a few model species.

Irrespective of their function and localization, all eukaryotic MutS homologues are encoded in the nuclear genome. But there is one exception. The mitochondrial genome (mtDNA) ^3^ of octocorals – a group of animals encompassing soft corals, blue corals, sea pens, and gorgonians – contains a MutS homologue (mtMutS)^4^ [20, 21]. Two hypotheses for the origin of mtMutS have been proposed. Earlier studies posited an intracellular transfer event of a nuclear MSH gene [22, 23]. More recent studies suggested a horizontal gene transfer (HGT)^5^ event, possibly from a giant virus [24, 25]. However, one critical piece of information was missing from these studies: none included cnidarian nuclear MSH genes. With the availability of genomic and transcriptomic data, it is now possible to analyze the changes in the nuclear MutS family that co-occurred with the acquisition of mtMutS.

Phylum Cnidaria, that contains the octocorals, is a group of mostly marine animals, with over 11,000 described species, comprising three major lineages: Anthozoa, Medusozoa, and Endocnidozoa. Anthozoa are sub-divided into Hexacorallia (sea anemones, hard corals, etc.), Octocorallia, and Ceriantharia (tube anemones). Medusozoa contains species from Cubozoa (box jellyfishes), Scyphozoa (true jellyfishes), Staurozoa (stalked jellyfishes), and Hydrozoa (hydroids, siphonophores, fire corals). Endocnidozoa consists of parasitic cnidarians from Myxozoa and Polypodiozoa [26, 27, 28].

Here, we analyzed the evolution of both nuclear-encoded and mitochondrial-encoded MutS homologues in Cnidaria, the only eukaryotic group containing MutS homologues encoded in both nuclear and organellar genomes. We also investigated MutS family composition in the rest of the animals.

## 2 Results

### 2.1 Characterization of MutS homologues in animals

To understand the origin and potential function of the octocoral mtMutS protein, we investigated the presence/absence of nuclear-encoded MutS homologs across Cnidaria. MSH2 and MSH6 were identified in all species studied, even in myxozoans that have been known to have experienced severe gene-loss and elevated rates of genome evolution [29] (Figure 1). The distribution of the other MSH proteins varied across different species. In particular, MSH3 was identified in only 11/17 species used in this study. MSH4 and MSH5 were recovered from 14/17 and 15/17 cnidarian species, respectively. By extending our analysis to 57 animal and 8 outgroup species, we found a similar pattern of conservation of MSH subunits across major animal phyla. While MSH2 and MSH6 were identified in nearly all the species analyzed, other MutS homologs were lacking in several animals. In particular, MSH3 was absent in majority of ecdysozoan and lophotrochozoan genomes analyzed in this study (Figure 4).

**Figure 1:**
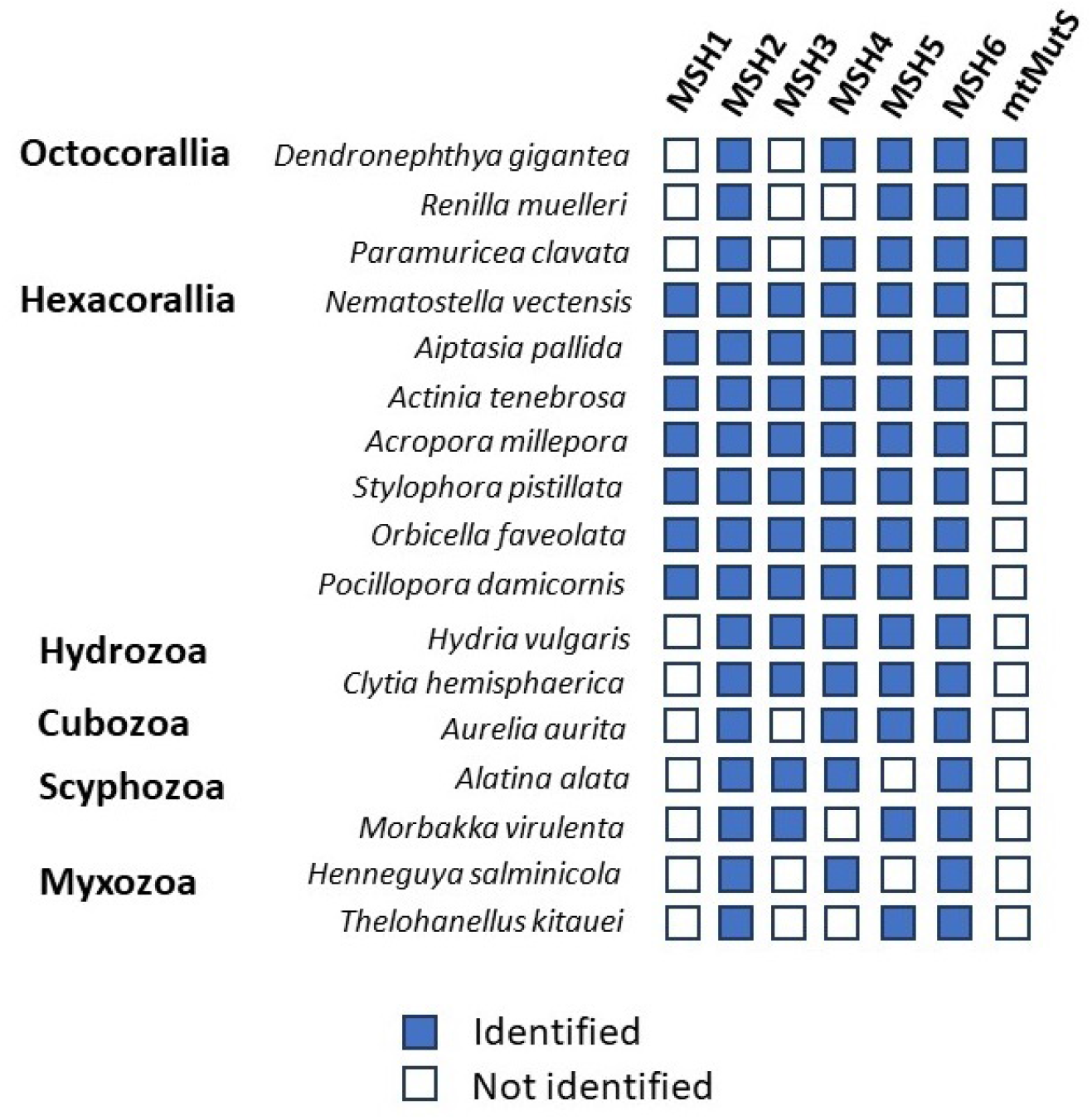
Characterization of nuclear and mitochondrial MutS homologues in cnidarian genomes.

### 2.2 Retention of MSH1 in several animal lineages

Surprisingly, we identified orthologues of fungal MSH1 in all hexacorals (phylum Cnidaria), several species of sponges (phylum Porifera), as well as *Trichoplax adhaerens* and *Hoilungia hongkongensis*, two species from phylum Placozoa. In each of the placozoan species, multiple MSH1 proteins were identified that resulted from Placozoa-specific duplication of the gene. MSH1 was also identified in choanoflagellates, the closest relatives of metazoans [30]. In contrast, we did not identify MSH1 in the genomes of octocorals, medusozoan cnidarians, ctenophores, and bilaterian animals.

We analyzed several key features of the opisthokont MSH1 proteins, including protein domain content, predicted overall protein structure, and the presence of a N-terminus MTS. Yeast MSH1 contains four protein domains: “MutS I”, “MutS II”, “MutS III”, and “MutS V”. The “MutS I” protein domain functions in DNA mismatch identification, and was found in all complete MSH1 proteins. A conserved phenylalanine within this domain critical for its function was found in all MSH1 proteins with the domain (Figure S1) (F77 in *Nematostella vectensis*) [31, 32]. All MSH1 proteins contained the “MutS V” (ATPase) domain. However, the “MutS II” and “MutS III” domains were not identified in four and one MSH1 proteins, respectively. An additional “MutS IV” domain was identified in 11 opisthokont MSH1 proteins, but not in yeast MSH1 (Figure 2).

**Figure 2:**
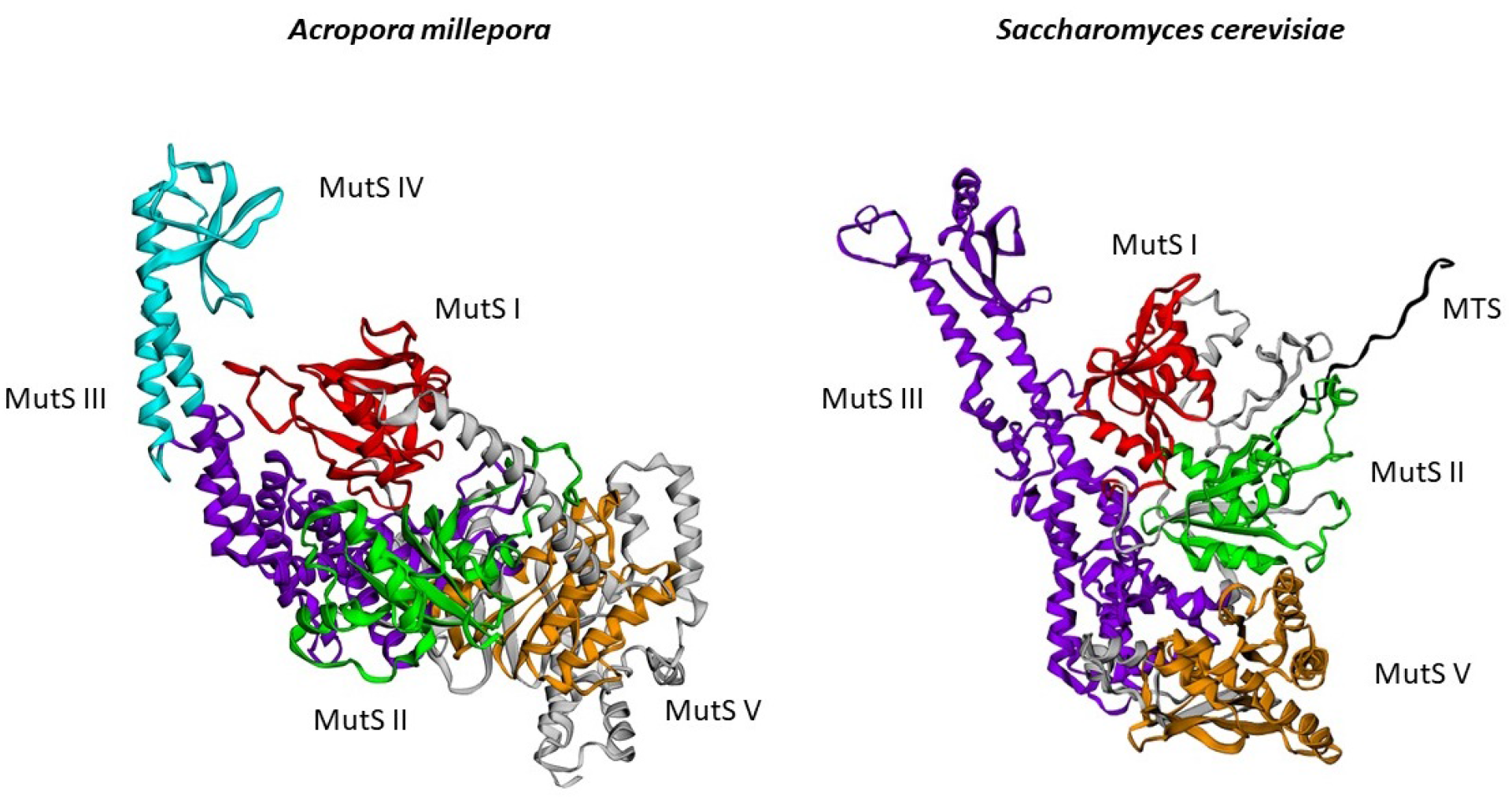
Protein structures of MSH1 from *Saccharomyces cerevisiae* and the hexacoral *Acropora millepora* as predicted by ROBETTA. Protein domains are indicated by different colors: MutS I (red), MutS II (green), MutS III (purple), MutS IV (blue), and MutS V (orange). The mitochondria-targeting signal in the fungal MSH1 is colored in black.

The yeast MSH1 contains a N-terminus mitochondria-targeting signal (MTS)^6^ that targets the protein to the mitochondria. DeepMito predicted that the majority of the MSH1 orthologues localized to the mitochondrial matrix (Supplementary Materials Folder S5). TargetP and MitoFates predicted that several non-bilaterian (5) and outgroup (3) MSH1 also possessed a MTS. However, none of the hexacoral MSH1 proteins were predicted to possess a canonical MTS.

### 2.3 Acquisition of mtMutS and loss of MSH1 in octocorals

While none of the octocoral genomes contained orthologues of yeast MSH1, all known octocoral mitochondrial genomes encode mtMutS. The analyzed mtMutS proteins were well-conserved with respect to size, hydrophobicity, and protein domain content (Supplementary Materials File S3). The size of octocoral mtMutS ranged from 957 amino acids in *Paragorgia* sp. 1075761 to 1,022 amino acids in *Briareum asbestinum* (mean=987 amino acids) (Figure S2). The mean GRAVY score of the mtMutS proteins, a measure of the hydrophobicity of the protein, was −0.11, and ranged from −0.046 in *Paragorgia* sp. 1075761 to −0.222 in *Calicogorgia granulosa* (Figure S3). The protein domain content of mtMutS was well-conserved. All mtMutS proteins contained the four domains: “MutS I”, “MutS III”, “MutS V”, and “HNH”. The mtMutS proteins from two species, *Briareum asbestinum* and *Sarcophyton trocheliophorum,* were predicted to contain a “MutS IV” domain, nested within the large “MutS III” domain.

The additional cnidarian MSH sequences identified in this study allowed us to analyze phylogenetic relationships between cnidarian mtMutS and MSH1-6 proteins. To better understand the origin of mtMutS, we built a phylogeny with MSH homologues, mtMutS, and blast hits to mtMutS and MSH1 (Figure 3). We saw that mtMutS is not related to MSH1, and is separated from all MSH subunits by a long branch. We identified a clade containing octocoral mtMutS and the MutS homologues from viruses and e-proteobacteria. We also noticed that there were some bacterial sequences closely related to MSH1. Hence, we searched for closest hits to all MSH homologues and found that most of them have closely related bacterial sequences (Figure S4).

**Figure 3:**
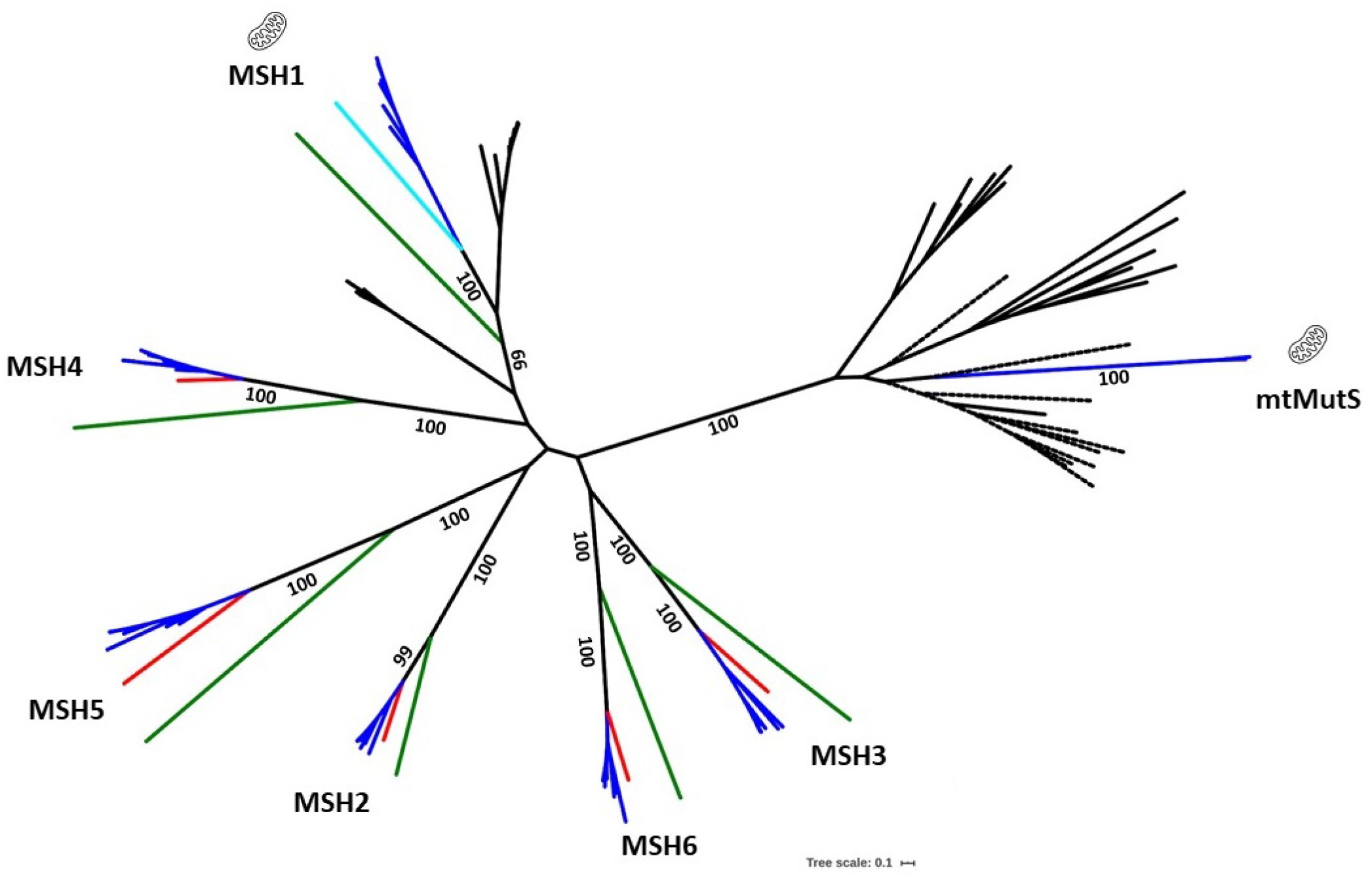
Phylogenetic tree of MutS homologues from cnidarians, bacteria, archaea, and viruses. Complete protein sequences were used to build this phylogeny using RAxML (1,000 boostraps). The branches are colored as follows: Cnidaria (blue), yeast (green), human (red), *Trichoplax adhaerens* (light blue), and prokaryotes (black). Dashed lines are used to depict viral species.

**Figure 4:**
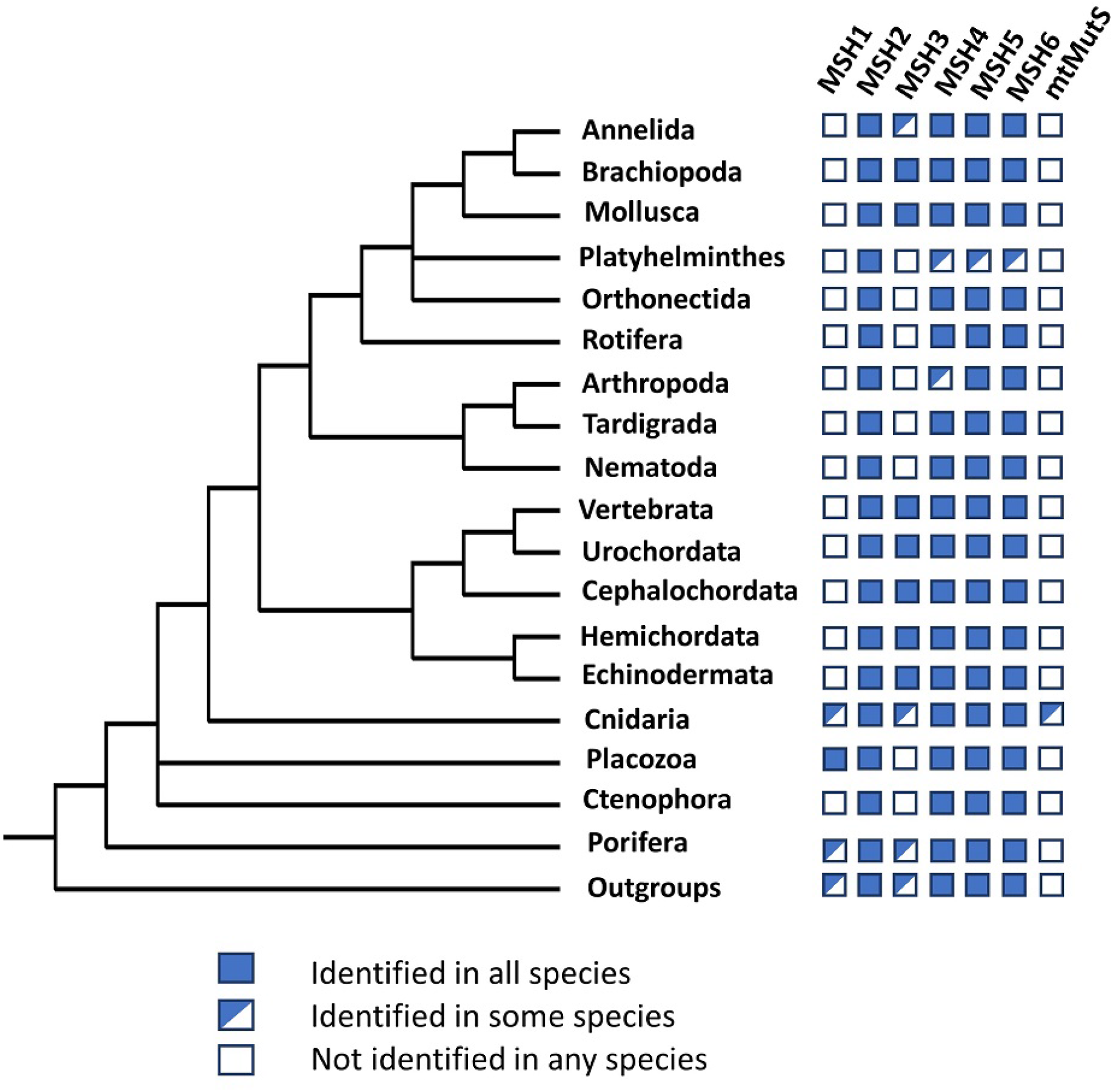
Distribution of MutS homologues across animal phyla.The phylogenetic tree used in this image was taken from [33]

## 3 Discussion

### 3.1 The evolution of the MSH family in animals

It is customary to say that MSH proteins are conserved across eukaryotes. Our study contradicts this notion and shows a spotty distribution of MSH3, MSH4, and MSH5 in animals. In particular, we were not able to identify MSH3 in several well-studied ecdysozoan and lophotrochozoan genomes by either orthology or HMMer-based approaches. The loss of MSH3 in animals has been reported in insects [34], including *Drosophila melanogaster* [35, 36]. Together with MSH2, MSH3 forms the MutS*β* heterodimer that recognizes longer insertions/deletions (up to 13 nucleotides) in mammals [8, 14]. MSH2 and MSH6 form the other heterodimer (MutS*α*) that identifies single-base mismatches, and smaller insertion/deletions (up to 3 nucleotides) [37, 38]. When both heterodimers are present, the loss of MutS*β* causes a weaker mutator phenotype compared to the loss of MutS*α* [39, 38]. However, when the MutS*β* heterodimer is lost, its function can be compensated by the MutS*α* heterodimer [36]. The remaining two MSH homologues with inferred losses in animals-MSH4 and MSH5, have roles in meiotic recombination instead of DNA repair [13]. These two proteins are known to be absent in the genome of *D. melanogaster*, where it has been proposed that other proteins have adopted their roles [35].

There is an additional complication in our understanding of MutS family evolution in eukaryotes. While some studies suggest that MSH proteins form a monophyletic clade and that their evolution can be explained by several rounds of duplication of an ancestral MutS homologue, most phylogenetic studies reconstruct eukaryotic MSH family as a paraphyletic group with some bacterial sequences closely related to MSH1 [25, 24]. Here, we found that most of the MSH sub-families had bacterial MutS sequences more closely related to them than sequences from other MSH subfamilies (Figure S4). Further studies are required to clarify whether this result indicates HGT of eukaryotic MSH gene to bacteria, multiple independent origins of eukaryotic MutS homologues, or contamination in the dataset.

### 3.2 Orthologues of yeast MSH1 in non-bilaterian animals

One of the surprising results of this study was the discovery of MSH1 orthologues in several phyla of non-bilaterian animals, as well as choanoflagellates. Although MSH1 was one of the first two MutS homologues found in eukaryotes, it was considered to be absent in animals [34]. By contrast, our study suggests that this gene was present in the common ancestor of all animals, and lost multiple times. Because yeast MSH1 is involved in mitochondrial DNA repair, we propose that a similar function of MSH1 was present early in animal evolution, and is still retained by some non-bilaterian animals. Interestingly, while both the original yeast MSH1 and the majority of MSH1 orthologues analyzed in this study were predicted to possess a MTS, the hexacoral MSH1 proteins lacked a MTS. However, this is not unexpected. In our previous analysis, we found that several non-bilaterian proteins with a high probability of mitochondrial localization lacked canonical MTS [40]. MSH1 is still predicted to be imported to mitochondria by DeepMito and likely uses an alternative targeting signal for mitochondrial import. While we found support for a clade that included opisthokont MSH1 and some eubacterial MutS proteins, we were unable to identify phylogenetic relationships within that clade, *i.e.* we were unable to conclude whether opisthokont MSH1 proteins were monophyletic.

### 3.3 The HGT origin of octocoral mtMutS

The identification of MSH1 in hexacorals and MSH2-6 from all major cnidarian groups allowed us to re-examine the phylogenetic relationships among the MutS homologues in Cnidaria. The evolutionary scenario proposed by Pont-Kingdon *et al.* (1998) [22] suggests that octocoral mtMutS resulted from the nucleus-to-mitochondria transfer of a MutS gene. In contrast, our results provide further support for the hypothesis that mtMutS was acquired via HGT from either a bacterium or a giant virus [25, 24]. The idea that giant viruses acted as vectors of mtMutS into octocoral mitochondria does have some additional support. First, HGT from giant viruses has likely happened one more time in the MutS family of proteins, with the transfer of pMSH1 in land plants [41]. Second, giant viruses are among the main groups of viruses found on the octocoral *Gorgonia ventalina* [42, 43, 44] and MutS orthologues from giant viruses are abundant in marine environments [24]. Finally, viral proteins have replaced mitochondrial proteins of eubacterial origin before, including RNA polymerase, DNA polymerase, and the primase-helicase TWINKLE [45]. However, mtMutS would be the first possible instance of HGT into the animal mitochondrial genome.

### 3.4 MSH1, mtMutS, and mitochondrial MMR in animals

Although we do not know whether hexacoral MSH1 and octocoral mtMutS are functionally similar, several lines of evidence support this hypothesis. Based on its orthology to yeast MSH1, conserved protein domain architectures in opisthokonts, and the inferred mitochondrial localization of opisthokont MSH1, animal MSH1 are likely to be involved in mitochondrial DNA repair. Similarly, octocoral mtMutS has conserved protein domain content, and functional residues suggesting a conserved function [25]. The distribution of MSH1 and mtMutS in animals could also provide insight into the function of the two proteins. Anthozoan mitochondrial genomes display some of the lowest rates of sequence evolution in animals, and also have either mtDNA-encoded or mitochondria-targeted MutS homologues [26]. Higher rates of mitochondrial sequence evolution are observed in medusozoan and endocnidozoan cnidarians, as well as ctenophores and bilaterian animals, which lack mitochondrial MutS homologues. Thus, we propose that both MSH1 and mtMutS are involved in mitochondrial DNA repair, and their loss correlates with higher rates of mitochondrial sequence evolution in species that have lost them. Additional research is being conducted by us and collaborators to characterize the function and structure of hexacoral MSH1 and octocoral mtMutS.

In conclusion, our study suggests that a viral MutS gene has integrated into the mitochondrial genome of octocorals via HGT and functionally replaced the MSH1 gene that was present in the common ancestor of Anthozoa. Furthermore, we demonstrate that the MSH1 gene is present in several phyla of non-bilaterian animals and that its loss correlates with higher rates of sequence evolution in the mitochondrial genomes of ctenophores and bilaterian animals.

## 4 Materials and Methods

### 4.1 Data acquisition

We downloaded the predicted proteomes of *Homo sapiens* (human), *Mus musculus* (mouse), *Saccharomyces cerevisiae* (yeast), and *Arabidopsis thaliana* (thale-cress) from UniProt [46]. These species are referred to as “reference species” because their MSH proteins have been well-characterized. To characterize MSH proteins in the phylum Cnidaria, we downloaded protein models from the genomes of 17 cnidarian species (10 anthozoan, 5 medusozoan, and 2 endocnidozoan species) (Supplementary Materials File S1). To identify MSH proteins across all major animal phyla, predicted proteins from the genomes of 57 animal species along with 8 non-animal outgroup species were downloaded from sources listed in Supplementary Materials File S2. For the analysis of the mtMutS protein in octocorals, we downloaded all publicly-available mitochondrial genomes of octocorals (89 species) from GenBank (Supplementary Materials File S1).

### 4.2 Identification of MutS homologues

#### 4.2.1 Phylum Cnidaria

The protein models from the 17 cnidarian species and the four reference species were used as input for OrthoFinder v2.4.0 to identify groups of orthologous proteins (OGs)^7^ in cnidarian and reference species (using default parameters) [47]. OGs containing MSH proteins from reference species were extracted for analysis. BlastP was used to remove potential contamination, and to identify homologues of the MSH proteins from these OGs in the Non-Redundant protein database (NR) (e-value:1e^-5^) [48, 49]. We retained only those proteins with a top BlastP hit from Cnidaria. Since some of the cnidarian MSH proteins were fragments, we created a manually-curated set of MSH proteins from cnidarians for downstream phylogenetic analysis (Supplementary Materials Folder S3).

We extracted the mtMutS gene from the 89 publicly-available octocoral mitochondrial genomes. For this, we re-annotated all mitochondrial genomes using MITOS2 [50]. We identified the MutS gene in all but one octocoral species-*Paragorgia* sp.USNM 1075769. The mtMutS gene from the latter species was riddled with multiple stop codons, possibly from mis-assembly of the mitochondrial genome. Hence, we excluded it from our dataset.

#### 4.2.2 Other species

OrthoFinder v2.4.0 was used to identify groups of orthologous proteins within the 57 animal and the 8 outgroup species (Supplementary Materials File S2). OGs containing MSH proteins from reference species were extracted for analysis. We used an HMMer-based approach to further examine instances of absence of MSH proteins in the genomes analyzed. For this, we downloaded reviewed sequences of MSH1-6 from Uniprot. Using these reviewed entries and MSH proteins identified in two well-studied cnidarians, *Nematostella vectensis* and *Hydra vulgaris,* we built HMM profiles of each MSH protein (MSH1-6) using HMMer v3.1.2 [51]. These profiles were then used to search for MSH proteins in the animal genomes (e-value:1e^-5^). For each animal genome, the results of the HMMer searches were downloaded, filtered based on sequence length (proteins below 500 amino acids in length were removed), and aligned to sequences of MSH1-6 from human and yeast using MAFFT v7 (“–auto” option) [52]. RAxML v8.2.11 was then used to build a phylogenetic tree for the resulting alignment with automatic selection of the substitution model and rapid bootstrapping with 100 resamples (“-m PROTGAMMAUTO -p 12345 -x 12345 -# 100”) [53]. The resulting phylogenetic trees were manually inspected to identify MSH subunits.

### 4.3 MutS characterization

Multiple tools were used to predict the sub-cellular localization of the MSH1 identified in this study: DeepMito [54], TargetP v2.0 [55], and MitoFates [56]. TargetP was used using the “Nonplant” option, DeepMito was used with default parameters, and MitoFates was used with the “Metazoa” option for animal sequences and “Fungi” option for the *S. cerevisiae* sequences. Protein domains were identified using the HMMer web-server [57]. Protein domain architectures were visualized using Illustrator of Biological Sequences (IBS) [58]. The 3D structure of MSH1 was predicted using the ROBETTA webserver [59] and visualized using EzMol [60]. GRAVY (Grand Average of Hydropathy) scores were calculated using (http://www.gravy-calculator.de/). The GRAVY score is a measure of the hydrophobicity of the protein, and is calculated as the sum of the hydropathy values of all amino acids divided by the length of the protein. Protein alignments were visualized by ESPirit v3 [61].

### 4.4 Phylogenetic analysis

BlastP was used to identify homologues of the hexacoral MSH1 and octocoral mtMutS proteins from the NR database. Using the octocoral mtMutS as a query, the top 10 best BlastP hits (based on e-value) from 1) Bacteria, 2) Archaea, 3) Viruses, and 4) Eukaryotes, were extracted from the NR database. Using the hexacoral MSH1 as query, the top 10 best BlastP hits from the NR database were downloaded (Supplementary Materials Folder S4). The manually-curated set of MutS homologues from cnidarians and the protein sequences resulting from the BlastP searches were aligned using MAFFT v7 (“–auto” option). TrimAI v1.2 was used to remove the poorly-aligned positions in the alignment (“automated1” option) [62]. The phylogenetic tree was reconstructed using RAxML v8.2.11 with automatic selection of the substitution model and rapid bootstrapping with 1,000 resamples (“-m PROTGAMMAUTO -p 12345 -x 12345 -# 1000”). RAxML identified the LG model as the best-scoring model of substitution. The resulting phylogenetic tree was visualized in iTOL v5.7 [63].

BlastP was used to identify the top 10 best BlastP hits (based on e-value) from prokaryotes, viruses, and eukaryotes, from the NR database for MSH1-6 from yeast and *Nematostella vectensis*. CD-HIT was used to cluster the sequences of the BlastP hits and the homologues of mtMutS identified from the NR database (described above) at 80% (“-c 0.8 -n 5”). These resulting sequences, along with MutS homologs from human, yeast, *N. vectensis*, and the octocoral *Dendronephthya gigantea* (MSH1-6 and mtMutS) were aligned using MAFFT (“–auto” option). TrimAI v1.2 was used to remove the poorly-aligned positions in the alignment (“automated1” option). RAxML v8.2.11 was used to construct the phylogenetic tree with automatic selection of the substitution model and rapid bootstrapping with 500 resamples (“-m PROTGAMMAUTO -p 12345 -x 12345 -# 500”).

### 4.5 Availability of the data

The sequences, scripts, and other supplementary materials from this study can be found on the project repository at Open Science Framework (https://osf.io/rn9ft/): Muthye V, Lavrov D (2020) Data for Dynamic evolution of the MutS family in animals: multiple losses of MSH paralogues and gain of a viral MutS homologue in octocorals. doi:10.17605/OSF.IO/RN9FT.

## 5 Acknowledgement and funding sources

We thank Dr. Cameron Mackereth and Dr. James Stewart for their comments on earlier version of the manuscript. This work was supported by a Research Grant from Human Frontiers Science Program (Ref.-No: RGP0050/2019)

## Supplementary Materials

**Figure S1:**
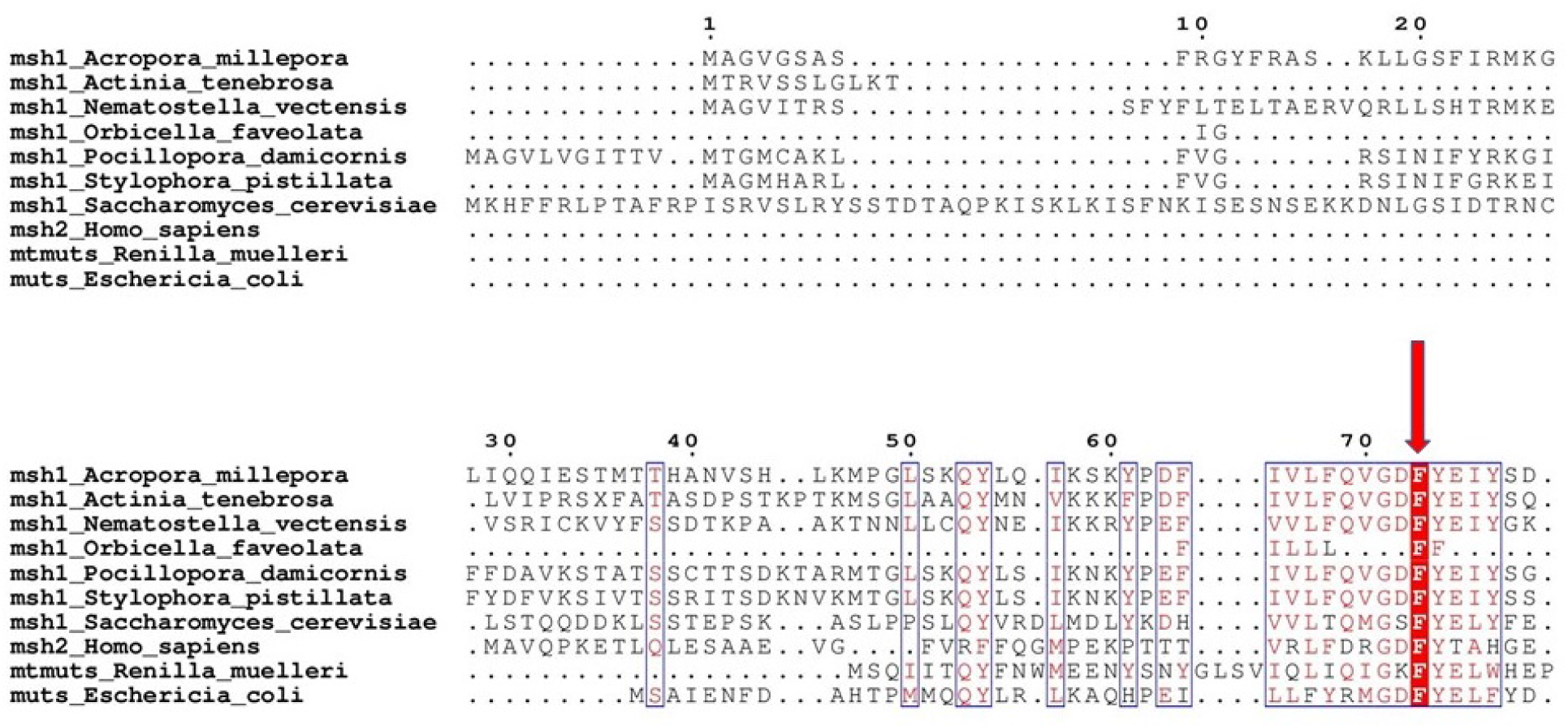
The N-terminus of the alignment of cnidarian MSH1 homologues along with mtMutS from *Renilla muelleri*, MSH2 from human, and MutS1 from *E. coli*. The conserved phenylalanine residue required for mismatch recognition in the domain “MutS I” is annotated with an arrow.

**Figure S2:**
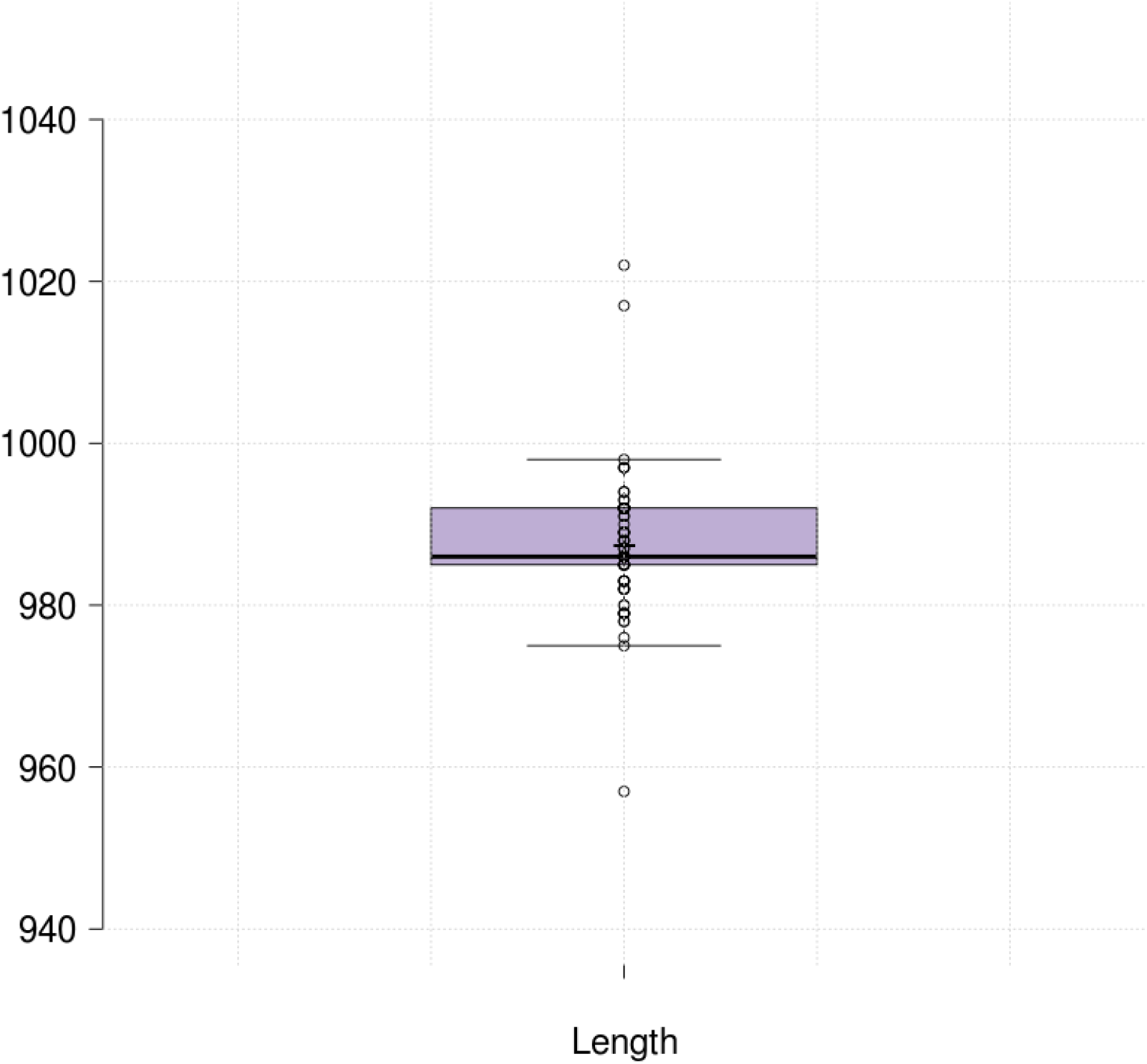
Box-plot depicting the distribution of protein lengths of octocoral mtMutS sequences (amino acids). The mean is denoted by a plus-sign (+) and the median is denoted by a horizontal bar (-).

**Figure S3:**
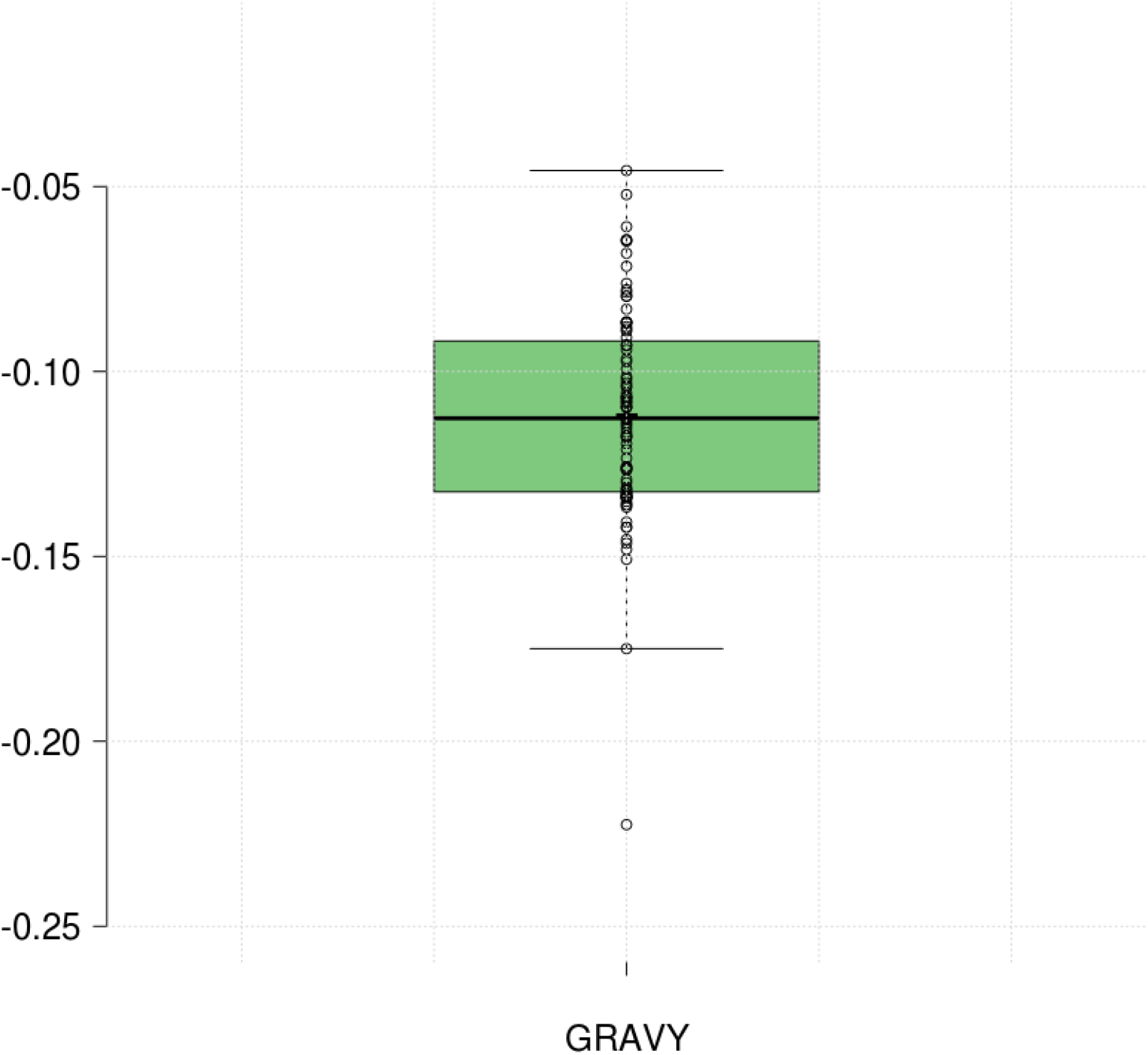
Box-plot depicting the distribution of GRAVY scores of octocoral mtMutS sequences. The mean is denoted by a plus-sign (+) and the median is denoted by a horizontal bar (-).

**Figure S4:**
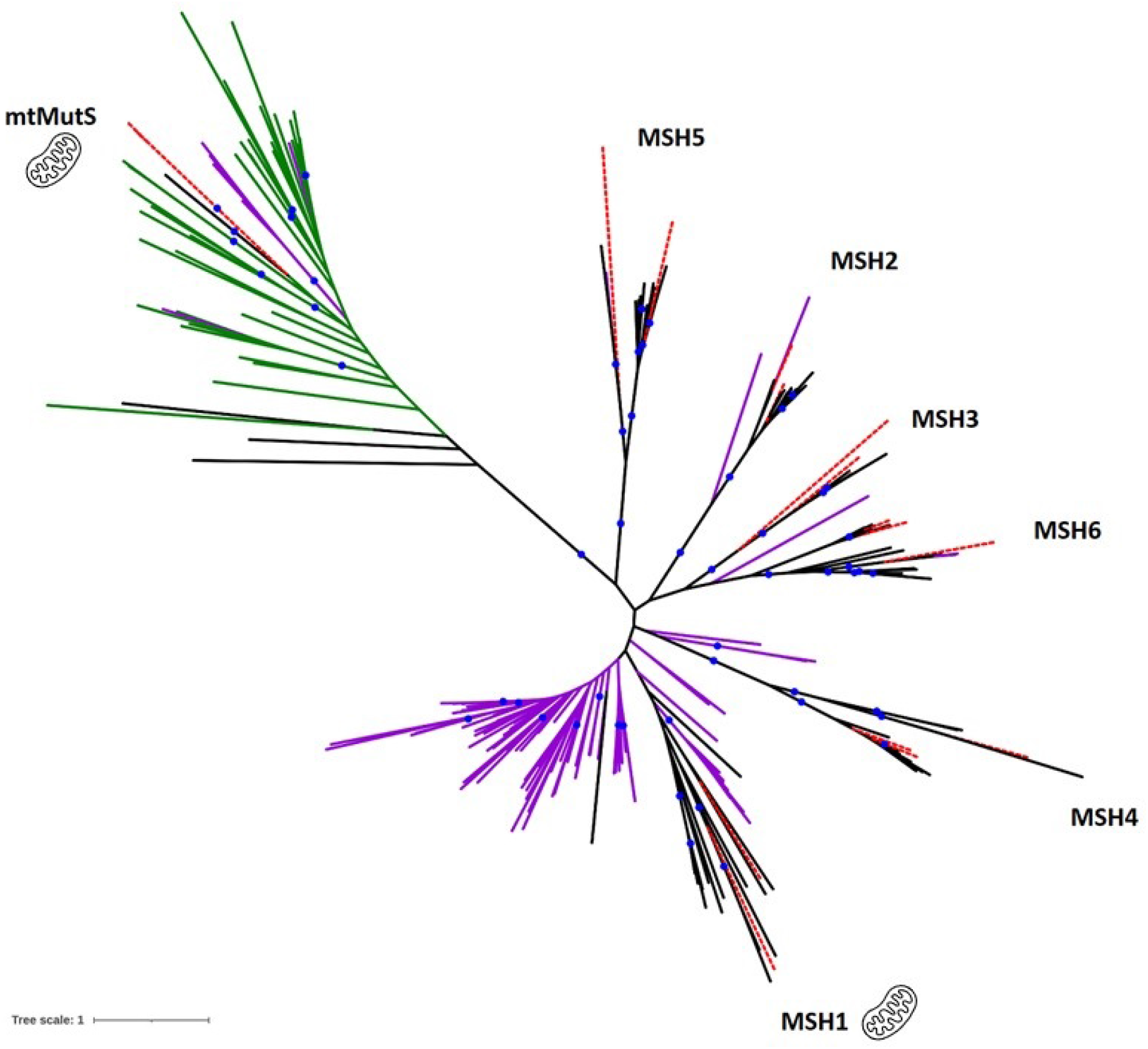
Phylogenetic tree of MutS homologues from eukaryotes, bacteria, archaea, and viruses. Complete protein sequences were used to build this phylogeny using RAxML (500 boostraps). The branches are colored as follows: MutS proteins from human, yeast, *Nematostella vectensis, Dendronephthya gigantea* (red dashed lines), viruses (green), prokaryotes (purple), and eukaryotes (black). Blue circles indicate bootstrap support values of at least 90%

## Footnotes

1 MMR: Mismatch Repair Pathway

2 pMSH1: plant-MSH1

3 mtDNA: mitochondrial genome

4 mtMutS: mitochondrial MutS

5 HGT: Horizontal Gene Transfer

6 MTS: mitochondria-targeting signal

7 OG: Group of Orthologous proteins

